# Circuit-specific gene editing for precision modulation of neuronal activity with CRISPR-rabies virus

**DOI:** 10.1101/2025.11.03.686433

**Authors:** Zijian Zhang, Jing Xu, Elizabeth A. Matthews, Samantha Deasy, Lewei He, Diego Arroyo, Enhui Pan, Arvin Deshmukh, Kiran Shehnaz Kaur, James O. McNamara, Derek G. Southwell, Charles A. Gersbach, Ru-Rong Ji, Dmitry Velmeshev

## Abstract

CRISPR gene editing has revolutionized our ability to study and manipulate specific genes, enabling novel insights into gene function and potential therapies for brain disorders. Recent advances in cell-type-specific regulatory elements and viral delivery systems have made precise *in vivo* gene editing possible. However, neurons with similar molecular profiles can belong to different circuits, complicating efforts to manipulate circuit function and behavior. To address this, we developed CRISPR-rabies virus (CRV), which leverages the trans-synaptic spread of rabies virus to enable gene editing within anatomically defined neural circuits. By pairing CRV with cell type-specific Cas9 expression, we achieved targeted gene modifications in specific circuits. We demonstrate that CRV can modulate sodium and potassium channel expression in parvalbumin interneurons, thereby effectively regulating synaptic transmission of pyramidal neurons in the CA3 region of the hippocampus. Its compatibility with 3′-capture single-cell RNA-seq allows simultaneous circuit perturbation and molecular profiling. In summary, CRV allows precise circuit-level gene modulation, providing a platform for studying gene function in neural circuits and developing novel gene therapies for brain disorders.

## Introduction

The adoption of the CRISPR–Cas9 system has revolutionized functional genomics and therapeutic development, expanding from DNA cleavage for gene disruption to versatile platforms for epigenetic modulation, base editing, and prime editing, with multiple clinical applications already in advanced stages^1–6^. In neuroscience, CRISPR has already transformed experimental possibilities. CRISPR interference (CRISPRi) and CRISPR activation (CRISPRa) allow precise control of gene expression without introducing DNA breaks, an especially valuable feature in post-mitotic neurons that lack robust DNA repair mechanisms^7–11^. With these tools, researchers can finely tune gene expression in individual neurons, manipulate genes within defined brain regions, and generate transgenic models that recapitulate human neurological disorders. However, a critical gap remains: the ability to manipulate genes at the level of specific neuronal circuits. Circuits are the organizing units through which the brain processes information, drives behavior, and shapes emotion; when they malfunction, neuropsychiatric and neurological disorders follow^12–14^. Despite its central importance, circuit-level gene editing remains out of reach: we lack delivery platforms that can genetically manipulate specific microcircuits without perturbing surrounding networks.

Realizing the full potential of CRISPR in neuroscience requires effective delivery strategies. Viral vectors have become key players for this purpose. For example, lentiviral vectors are widely used for *ex vivo* editing of dividing cells, while adeno-associated virus (AAV) is the dominant choice *in vivo* due to its favorable safety profile and non-integrating life cycle^15,16^. Recent advances in AAV engineering by incorporating cell type–specific regulatory elements and designing serotypes like AAV-PHP.eB that cross the blood–brain barrier in mice, have expanded the scope of genetic interventions^17^. Nevertheless, current AAV-based approaches remain fundamentally limited in their ability to achieve widespread, circuit-specific gene editing in the brain. This is a key requirement for probing complex neuronal functions and developing therapies for neurological disease. Also, neuronal transcriptomic identity does not fully predict connectivity, as neurons of the same molecular type can form distinct patterns of synaptic input and output^18,19^.

Monosynaptic rabies virus (RV) has long been the gold standard for mapping direct synaptic inputs, spreading from genetically defined “starter cells” to their presynaptic partners^20–22^. In this study, we engineered a CRISPR-rabies virus (CRV) that uses endogenous tRNA processing to generate functional sgRNAs from rabies transcripts. By pairing CRV with cell–type–specific Cas9 expression, we achieved targeted gene modifications within defined microcircuits. As a proof of principle, we demonstrate that CRV-mediated perturbation of sodium and potassium channels in parvalbumin interneurons effectively modulates pyramidal neuron firing in the hippocampal CA3, establishing direct causal links between molecular regulation and circuit dynamics. Furthermore, combining CRV with single-cell sequencing allows circuit-specific gene perturbation and comprehensive molecular characterization. Together, these findings establish CRV as a CRISPR delivery tool and a circuit-resolved functional genomics platform. enabling precise dissection of gene function in neural circuits and providing a foundation for circuit-aware therapeutic strategies for brain disorders.

## Results

### Development of CRV for sgRNA delivery

To enable circuit-specific gene editing, we first sought a viral vector capable of transsynaptic spread. Since monosynaptic rabies virus is a widely adopted tool for transsynaptic tracing^20–23^ that allows carrying gene cargoes between synaptically coupled cells, we chose this viral vector as the vehicle for delivering sgRNA. However, unlike widely adopted gene delivery vectors such as AAV or lentivirus, rabies virus is a negative-sense single-stranded RNA virus^24,25^, and its genome organization and transcription mechanism pose unique challenges for sgRNA expression. Specifically, rabies uses a promoter sequence at the 3’ end of its genome to initiate transcription and uses its own viral RNA polymerase, the viral L protein^24,26^. Consequently, traditional approaches for small RNA expression, such as driving sgRNA transcription from a U6 promoter via host RNA polymerase III^24,26^, are ineffective. Moreover, transcripts generated by the viral L protein carry a 5’ cap and 3’ poly(A) tail, which could interfere with the proper structuring of the sgRNA and its binding to Cas proteins^27–29^. We circumvented this obstacle by encoding a tandem cassette of Transcription On-tRNA-sgRNA-tRNA-sgRNA-tRNA-Off into the genome of B19 strain of rabies virus (**Fig. 1a**). Since tRNA is cleaved by RNase P and Z in the nucleus to precisely excise tRNA, this process also releases sgRNA without 5’ and 3’ redundant sequences^27,30,31^.

**Figure 1.**
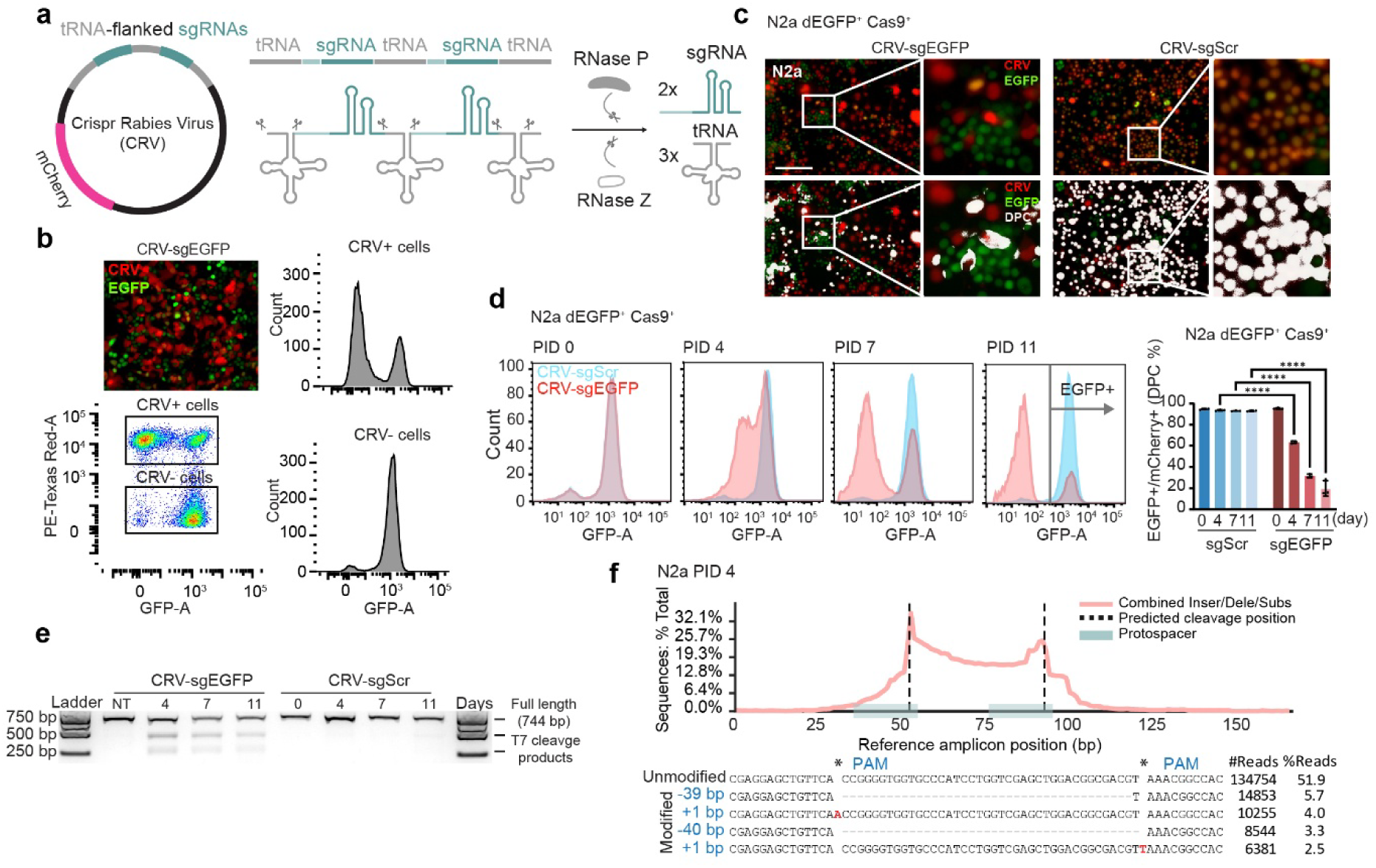
tRNA processing enables functional sgRNA delivery by rabies virus. **a)** Design of tRNA-sgRNA cassette for expression of sgRNAs by rabies virus. **b)** Microscopy and flow cytometry analyses showing loss of EGFP expression in cells infected with CRV (scale bar, 150 μm). **c)** N2a-dEGFP-Cas9 cells infected with CRV expressing nuclear-localized mCherry and carrying either an *EGFP* targeting sgRNA or a scrambled control. Double-positive cells (DPCs) are highlighted in white. Data are presented as mean ± s.d. Two-way ANOVA with multiple comparisons (****p < 0.0001; ns, not significant). **d)** Flow cytometry analysis of EGFP signal from day 0 to 11 in mCherry⁺ cells following CRV infection. **e)** T7E1 assay on the EGFP locus amplified 0-11 days after infection at MOI = 1. **f)** CRISPResso2 analysis of mutation position distribution at the EGFP target locus in 293T cells from e) infected with CRV for 4 days.

To test the functionality of the CRV as an sgRNA delivery vector, we designed two GFP-targeting sgRNAs simultaneously from a dual-sgRNA cassette, and co-expressed an mCherry fluorescent protein reporter. CRV particles were produced and applied to human embryonic kidney 293T/17 (293T) and mouse neuroblastoma Neuro-2a (N2a) cell lines, stably expressing destabilized EGFP (dEGFP) and Cas9. CRV infection led to the emergence of an EGFP negative cell population among mCherry⁺ (CRV+) cells, whereas nearly all uninfected cells (CRV-) remained EGFP positive **(Fig. 1b)**. Over the course of infection, mCherry and EGFP signals became progressively mutually exclusive, indicating successful EGFP editing in infected cells **(Fig. 1c and s1a)**. At a multiplicity of infection (MOI) of 0.3, we observed a progressive decrease in the proportion of EGFP⁺ cells among the CRV-infected (mCherry⁺) population over time, reaching a 50–80% reduction by post-infection day (PID) 11 in N2a and 293T cells (**Fig. 1d and s1b**), respectively. T7 endonuclease I (T7E1) assay (**Fig. 1e and s1c**). To assess editing outcomes, we designed an amplicon spanning both predicted Cas9 cleavage sites, enabling detection of both individual indels at each site as well as deletions between the two cut sites. Deep amplicon sequencing at day 4 post-infection confirmed robust editing at both target sites, and revealed frequent deletion events spanning the intervening region, consistent with dual-cut outcomes expected from simultaneous sgRNA expression (**Fig. 1f and s1d**). In summary, we established and validated a rabies virus–based gene-editing vector capable of delivering sgRNAs for precise reporter gene targeting.

### Comparison of CRV, lentivirus, and AAV in gene editing performance

After demonstrating that CRV can deliver sgRNAs and mediate *EGFP* disruption in the presence of Cas9, we next compared its gene-editing efficiency with two well-established viral vectors: AAV and lentivirus. All three vectors were engineered to express the same pair of sgRNAs targeting 2 sites of *EGFP* locus; AAV and lentivirus utilized the commonly used U6 promoter for sgRNA expression, while CRV employed tRNA-flanked sgRNAs driven by viral promoter. We infected fewer than 30% of cells with each virus, as measured by mCherry signal at PID 3. Subsequent analyses, including imaging, FACS, T7E1 assay, and amplicon NGS, were performed to assess editing efficiency (**Fig. 2a**). FACS analysis showed that CRV-mediated disruption of destabilized EGFP was slower than that of lentivirus in 293T cells (**Fig. s2a-c**), but in the neuronal cell line N2a, CRV achieved editing efficiency comparable to lentivirus at PID 7 (51% for CRV vs 50% for Lentivirus) (**Fig. 2b&c**). In both cell types, CRV significantly outperformed AAV, which exhibited much lower gene-editing efficiency under the same *in vitro* conditions (**Fig. 2b, c and s2a-c**). On day 4, infected cells were sorted by FACS and subjected to genomic DNA amplification. Amplicons were analyzed either by the T7E1 assay or by next-generation sequencing (NGS), followed by CRISPResso2 analysis (**Fig. 2d, e, and s2d, e**). These assays revealed efficient target editing, demonstrating that CRV can effectively induce mutations at levels comparable to those achieved with well-established AAV or lentiviral sgRNA delivery vectors.

**Figure 2.**
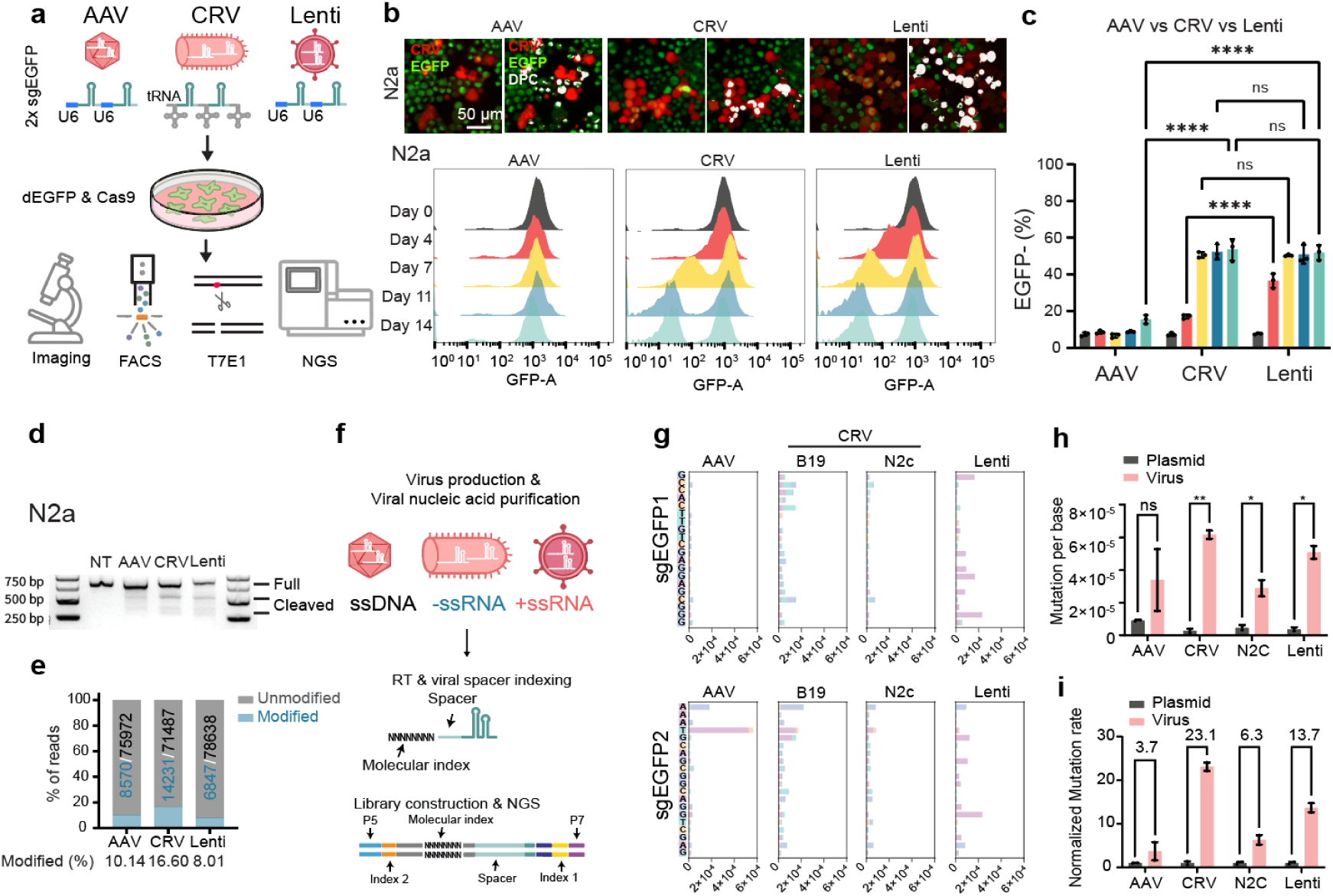
Comparison of efficiency and reliability between CRV, AAV, and lentiviral sgRNA delivery systems. **a)** Schematic overview of the comparative workflow and downstream readouts (microscopy and longitudinal flow cytometry of EGFP loss, T7E1 at the EGFP locus, and NGS of indexed amplicons). **b)** AAV, CRV, and lentivirus expressing mCherry together with two identical sgRNAs targeting EGFP. At day 11 post-infection, red (mCherry⁺, infected) N2a-dEGFP-Cas9 cells losing green fluorescence indicate successful targeting. Double-positive cells (DPCs) are highlighted in white. **c)** Flow cytometry analysis of EGFP expression from day 0 to 14 in mCherry⁺ cells following AAV, CRV, or lentiviral infection. **d)** T7E1 assay of the EGFP locus amplified from genomic DNA of mCherry⁺ cells, FACS-sorted 4 days after infection at MOI = 0.3 (CRV, Lenti) or ∼100,000 (AAV). **e)** Bar plot showing mutation frequency determined by NGS of adapter-ligated PCR products from (d). **f)** Schematic illustration of NGS-based analysis to quantify mutation frequency and distribution within gRNA sequences in viral genomes. **g)** Mutation frequency and types for sgEGFP1 and sgEGFP2. **h)** Bar plot comparing mutation frequencies (reads ≥ 3). Unpaired t-test. Data are presented as mean ± s.e.m. **i)** Bar plot showing mutation frequencies in the different viral sgRNA-encoding plasmids used for viral production (sgRNA expression plasmids) and in the corresponding viral genomes (reads ≥ 3). Multiple unpaired t-test. Data are presented as mean ± s.e.m.

### Assessment of the reliability of sgRNA delivery by CRV

A key requirement for any CRISPR-based tool is reliable and accurate delivery of sgRNA sequences, as mutations can compromise targeting specificity and efficacy. Unlike AAV or lentivirus, rabies virus production involves a prolonged recovery process, often requiring several weeks and multiple rounds of viral replication and amplification^32^. These steps increase the risk of introducing mutations^33–35^. To assess the fidelity of guide sequences delivery by CRV, we adopted a one-step recovery method to minimize replication cycles during virus production^34^. AAV and lentiviral vectors were produced and purified by standard protocol, through cesium chloride or sucrose ultracentrifugation, respectively. CRV particles were recovered and pseudotyped in BHK-EnvA cells, followed by ultracentrifugation. Viral genomes were extracted, and 12-nt random barcodes were incorporated during first-strand synthesis to correct amplification-induced mutations from sequencing artifacts (**Fig. 2f**). Amplicons were sequenced and analyzed to quantify mutation rates in the guide sequences and adjacent regions. Our results showed that the B19 strain of CRV exhibited a higher mutation frequency compared to other systems, and B19 showed around 23-fold higher mutation frequency than the plasmids used for its production (**Fig. 2g-I and s2f**). We also evaluated the rabies virus strain CVS-N2c (N2c hereafter), which has been widely used in neural circuit tracing. Compared to the more commonly used SAD-B19 strain, N2c exhibits lower cytotoxicity and is considered more attenuated, allowing better neuronal survival in tracing experiments. In addition, N2c retains efficient trans-synaptic spread capability and shows higher tracing efficiency than B19^36,37^. N2c strain demonstrated even lower mutation rates than AAV and lentivirus (**Fig. 2g-i**). Importantly, even in CRV-B19, which exhibited the highest mutation rate (6×10^-5^ per nt), approximately 99.9% of 20-nt sgRNA sequences remained mutation-free, underscoring the high fidelity of CRV as an sgRNA delivery system.

### Optimization of the efficiency and applicability of CRV

To enhance the applicability of CRV, we sought to optimize one of the major limitations of rabies virus, its inherent toxicity^23,38,39^, by reducing cytotoxicity while maintaining or improving gene-editing efficiency. To address this issue, we turned to the N2c strain due to its lower toxicity. N2c exhibits comparatively lower transgene expression per infected neuron than SAD-B19 in standard ΔG tracing contexts^36,37^, which likely limits sgRNA abundance and compromises editing efficiency (**Fig. 3a**). This observation aligns with prior reports that transgene/RNA expression level critically shapes both toxicity and functional readouts in engineered rabies vectors^36,40,41^. To restore editing efficiency, we explored two strategies. The first leveraged the ability of tRNAs to multiplex sgRNAs (**Fig. s3a&b**), thereby increasing sgRNA copy number. The second sought to enhance sgRNA processing by directing rabies RNA into the nucleus (**Fig. s3c**), where RNase P and RNase Z are active^42,43^. For the first strategy, we generated tandem tRNA–sgRNA–tRNA cassettes by PCR amplification and inserted them into the N2c genome using Gibson assembly. After transformation, we performed colony PCR and Sanger sequencing to identify successfully edited constructs containing different copy numbers of the tRNA–sgRNA units. Four to six-copy multi-sgRNA CRV-N2c achieved EGFP editing efficiencies comparable to CRV-B19 carrying two sgRNAs (**Fig. 3b**). Moreover, N2c exhibited a lower mutation rate than B19, comparable to AAV and lentivirus (**Fig. 2g-i**). This design thus balances reduced toxicity with robust editing, making CRV more suitable for *in vivo* circuit-level applications. The second strategy uses MS2-MCP, PP7-PCP, or BoxB-N22 systems^44,45^ with the RNA aptamer-targeting protein component fused to mCherry-NLS (**Fig. s3c**) and enriching rabies RNA in the nucleus. This approach successfully redirected RNA to the nucleus but did not improve the editing efficiency of CRV-B19 (**Fig. s3d, e**). This design may have sequestered essential rabies RNAs in the nucleus, whereas the rabies virus life cycle normally occurs in the cytoplasm^24,46^, thereby restricting its normal expression. Therefore, we adopted the route of multiplexing sgRNA in the CRV-N2c and utilized 4xsgRNA CRV-N2c in the rest of the experiments.

**Figure 3.**
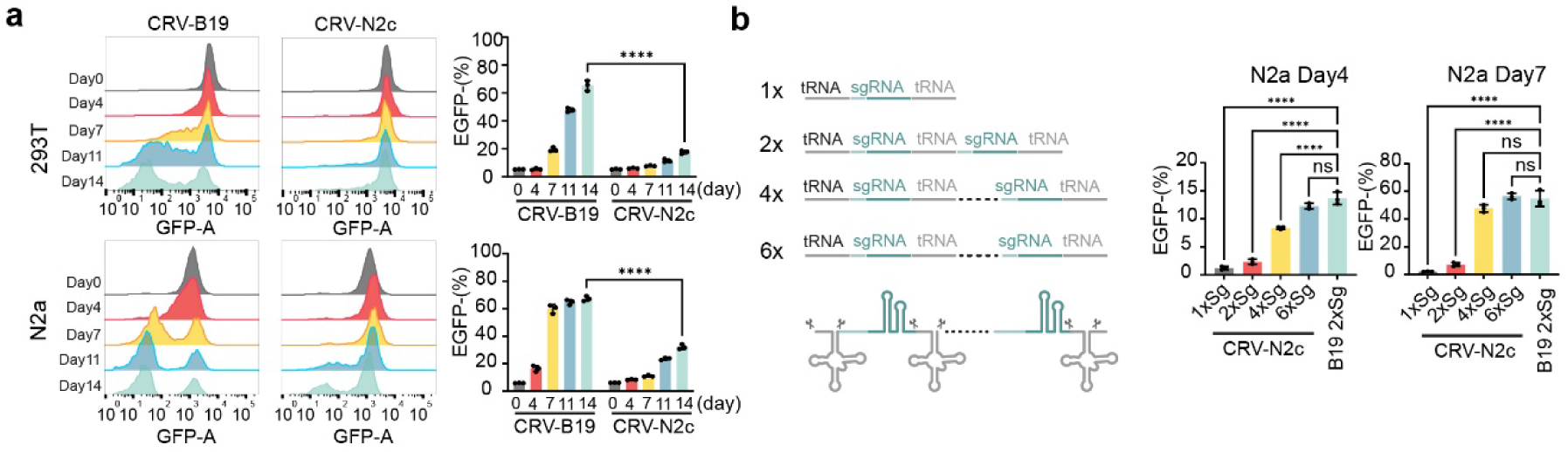
Multiplexed tRNA–sgRNA design enables efficient delivery by the attenuated rabies strain N2c. **a)** Flow cytometry analysis of EGFP loss from day 0 to 14 comparing CRV B19 and CRV N2c in 293T and N2a reporter cell lines. **b)** Flow cytometry analysis of N2c-CRV carrying different copy numbers of sgRNAs, compared with first-generation CRV based on the B19 rabies strain.

### CRV enables endogenous gene regulation

Having validated and optimized CRV’s performance in EGFP reporter systems, we next assessed its ability to modulate endogenous genes. We constructed CRV-N2c vectors carrying sgRNAs directed against the mouse *Scn1a* and *Kcna1* gene bodies or upstream regulatory regions^9,10^. To mimic transsynaptic gene editing, we developed an *in vitro* co-culture model in which CRV replicates in one cell type and infects another cell type. Specifically, B7GG cells^32^ were first infected with CRV (mimicking “postsynaptic” cells). After 24 hours, cells were washed and re-seeded into new plates, then co-cultured with N2a cells expressing Cas9 and EBFP (**Fig. 4a**). After 5 days of co-culture, mCherry⁺/EBFP⁺ cells (mimicking “presynaptic” cells) were isolated by FACS. T7E1 assays confirmed the presence of CRISPR-induced mutations at the targeted *Scn1a* and *Kcna1* loci (**Fig. 4b**). We next examined whether CRV is compatible with CRISPRa and CRISPRi, dCas-based systems frequently used to modulate gene expression. In mouse embryonic carcinoma P19 and neuroblastoma N2a cell lines, we tested two CRISPRa systems (dCas9-p300^47,48^ and dCas9-VPR^49^) and one CRISPRi system (dCas9-KRAB^50^). Both mRNA and protein analyses revealed significant changes in the expression of corresponding targets, confirming that these tools can effectively modulate endogenous gene expression (**Fig. 4c, d and s4a, b**).

**Figure 4.**
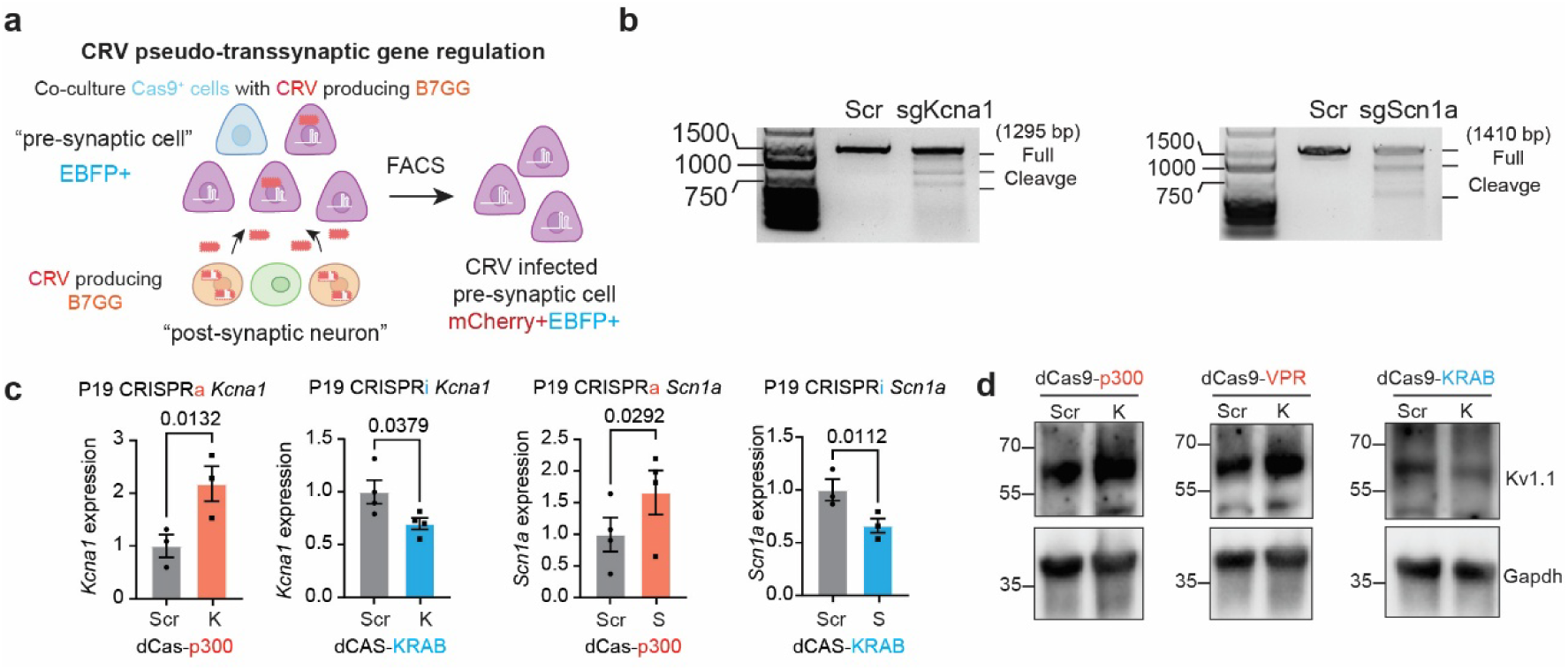
CRV targets endogenous genes and are compatible with CRISPRa/CRISPRi tools. **a)** Schematic of the in vitro co-culture model simulating trans-synaptic gene regulation by CRV. **b)** T7E1 assay confirming the efficiency of sgRNAs targeting the endogenous Scn1a and Kcna1 loci. **c-d)** CRV-mediated CRISPRi and CRISPRa regulation of gene expression, demonstrated by qRT-PCR (mRNA levels of Scn1a and Kcna1) and Western blot (protein levels Nav1.1 and Kv1.1). (Scr, scramble control; K, sgKcna1; S, sgScn1a). Unpaired t-test. Data are presented as mean ± s.e.m.

### CRV manipulates gene expression in defined neural circuits *in vivo*

To evaluate whether CRV-N2c can modulate gene expression within intact neural circuits, we used mice generated by crossing PV-Cre with Rosa26-LSL-dCas9-p300 (hereafter referred to as PV-p300) enabling selective expression of dCas9-p300 in parvalbumin (PV) interneurons **(Fig. s5a**). We injected AAV9-Camk2α-TVA-EGFP-oG into the hippocampal CA3 region to express the TVA receptor, optimized rabies glycoprotein (oG)^38^, and EGFP in pyramidal neurons, which served as starter cells for CRV infection and replication **(Fig. 5a and s5b**). One to two weeks later, CRV-N2c carrying sgRNAs targeting either *Kcna1* or *EGFP*, along with H2B-mPlum, was injected into the same region (**Fig. s5c, d**). To assess the effects of gene targeting, brain slices were collected two weeks post-CRV injection (**Fig. 5a**). A subset was processed for immunofluorescence staining to assess Kv1.1 protein levels (**Fig. 5b and s5e**). To evaluate downstream circuit effects, we prepared acute brain slices, and performed whole-cell patch-clamp recording of spontaneous inhibitory postsynaptic currents (sIPSCs) and excitatory postsynaptic currents (sEPSCs) in hippocampal CA3 from EGFP^+^&H2B-mPlum⁺pyramidal neurons (**Fig. 5c**). We observed that upregulating Kv1.1 in PV neurons led to a significant decrease in sIPSC frequency in the starter pyramidal neurons compare to sgEGFP group (p=0.0364, **Fig. 5d, e**). This result suggested a reduced inhibitory input stemming from decreased activity of PV neurons and a predicted disinhibitory effect on excitatory neurons. Furthermore, an increased sEPSCs frequency was found in pyramidal neurons after Kv1.1 upregulation in PVs (**Fig. 5f, g**). Conversely, when we upregulated Nav1.1 (a sodium channel that controls excitability) in presynaptic PV neurons, we observed the opposite effects: sIPSC frequencies in starter pyramidal neurons were increased and sEPSC frequencies showed a downward trend (**Fig. 5f, g**). This aligns with data from a transgenic mouse model, in which Nav1.1 is disturbed in PV neurons; there, pyramidal neuron hyperexcitability is observed, and the restoration of inhibitory tone suppresses associated seizures^9,51^. Taken together, these results validate that CRV can modulate specific genes in defined neuronal circuits and produce highly specific and predictable effects on both inhibitory and excitatory synaptic transmission of specific neuronal subtypes within a defined circuit.

**Figure 5.**
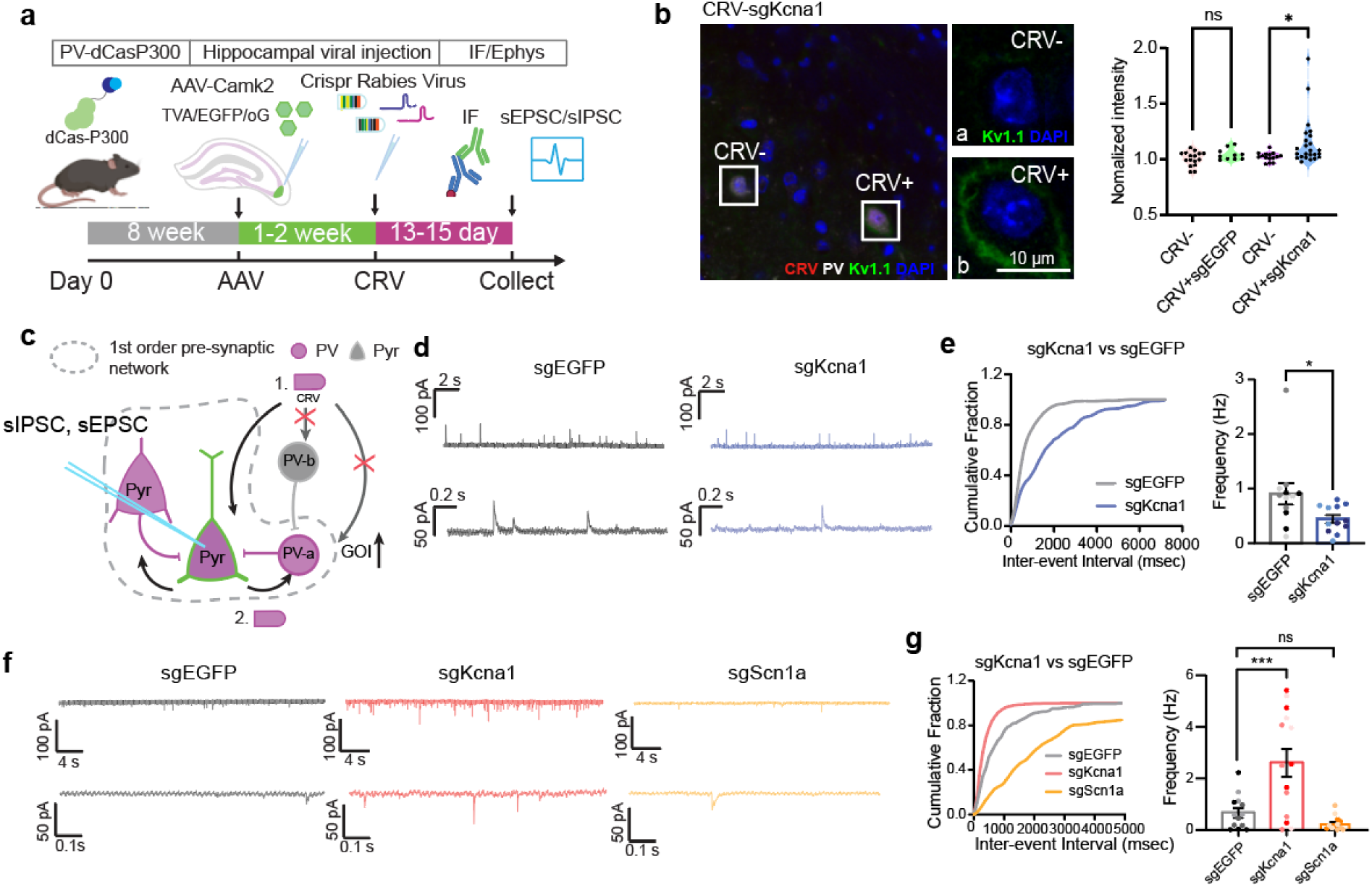
*In vivo* CRV-mediated circuit specific targeting of PV neuron alters postsynaptic sEPSCs and sIPSCs. **a)** Schematic of the *in vivo* experiment targeting the Kcna1 locus in PV neurons presynaptic to hippocampal CA3 pyramidal neurons. PV-p300 mice are injected with AAV9-Camk2-TVA-EGFP-oG to create Camk2a⁺ excitatory starter cells. Immunofluorescence detecting Kv1.1 expression in PV neurons, and whole-cell recordings measuring sEPSCs and sIPSCs in pyramidal neurons. **b)** Immunofluorescence analysis of Kv1.1 expression in PV neurons infected with CRV-sgEGFP or CRV-sgKcna1, and compared to uninfected cells in each group. Unpaired t-test (*p < 0.05; ns, not significant). **c)** Schematic of CRV-mediated trans-synaptic delivery enabling circuit-specific gene regulation in PV neurons. **d–e)** Representative sIPSCs traces **(d)** and quantification **(c)** of hippocampus pyramidal neurons recording showing the decreased sIPSC frequency after CRV-mediated activation of Kv1.1 expression in presynaptic PV interneurons (n =12 cells from 3 mice per group). Unpaired t-test (*p < 0.05). Data are presented as mean ± s.e.m. **f-g)** Respectively sEPSCs traces **(f)** and quantification **(g)** of hippocampus pyramidal neurons recording in three different groups (n =13, 14, 12 cells from 3 mice per group). One-way ANOVA followed by Dunnett’s multiple comparisons test (****p < 0.001; ns, not significant). Data are presented as mean ± s.e.m.

### Integration of CRV with single-cell sequencing for circuit-level gene function analysis

The combination of CRISPR-based perturbation with single-cell transcriptomics has emerged as a powerful approach to dissect gene function with cell-type and state resolution. This allows linking gene perturbations to cell-type–specific transcriptional responses with high cellular and gene throughput. However, most CRISPR systems express sgRNAs under a U6 promoter, resulting in non-polyadenylated transcripts, whereas most single-cell sequencing platforms rely on capturing polyadenylated RNA molecules^52–54^. Overcoming this limitation requires specialized assays to capture sgRNAs in single cells^55,56^. CRV offers a unique advantage: the initial sgRNA– tRNA transcripts expressed by CRV retain a 3’ polyadenylated tail prior to tRNA processing, making them directly compatible with standard 3’-based single-cell RNA sequencing platforms (e.g., 10x Genomics Chromium) (**Fig. 6a**). This inherent compatibility greatly simplifies CRISPR– scRNA-seq integration and reduces associated costs. To demonstrate this, we injected AAV9-Camk-TVA-EGFP-oG and CRV-N2c into the hippocampus (**Fig. 6a**), as in the experiments described above (**Fig. 5a**). Dissected tissue was used to isolate nuclei for fluorescence-activated nuclei sorting (FANS) enrichment of infected neurons based on H2B-mPlum fluorescence from three groups of mice: CRV-sgKcna1, CRV-sgScn1a, and control CRV-sgEGFP. Nuclei from the CRV-sgKcna1 group were processed separately, whereas sgScn1a and sgEGFP were pooled to test the feasibility of demultiplexing based on sgRNA sequences (**Fig. s6a**). Clusters and UMAP embeddings were generated with Seurat v5^57^, and major cell types were annotated using ScType markers^58^ (**Fig. 6b**). CRV (mPlum) exhibited substantially higher viral transgene RNA abundance compared with AAV (oG) under the Camk2 promoter, reaching 8.75 ± 1.25-fold of oG (mean ± SEM), consistent with our observation of stronger CRV RNA output and higher editing efficiency, and with prior reports that rabies genomes achieve high transcriptional levels^24,59^. Along the CRV genome, gene expression followed the expected N to L decreasing gradient (**Fig. 6c**), matching the well-described promoter-proximal attenuation of nonsegmented negative-strand RNA viruses^25,60^. Within the hippocampus, CRV-infected nuclei from CA3 showed the highest viral RNA levels—higher than other regions—which is consistent with CA3 being the injection locus (earlier infection time) whereas cells in other regions are likely presynaptic partners labeled later. Gad1 and Pvalb expression were used to identify PV interneurons (**Fig. 6d, e**). Notably, cells expressing their respective sgRNAs exhibited clear up-regulation of the intended target gene in PV interneurons (**Fig. 6f**), but not in other interneuron subtypes (e.g., VIP or SST) (**Fig. s6b, c**); closely related paralogs such as *Scn2a* and *Kcna2* remained unchanged (**Fig. 6g**), and canonical markers (*Gad1*, *Syn1*) showed no global shifts. These results demonstrate the specificity of CRV in regulating gene expression and establish a new Perturb-seq platform that enables circuit-informed functional genomics in the mammalian brain.

**Figure 6.**
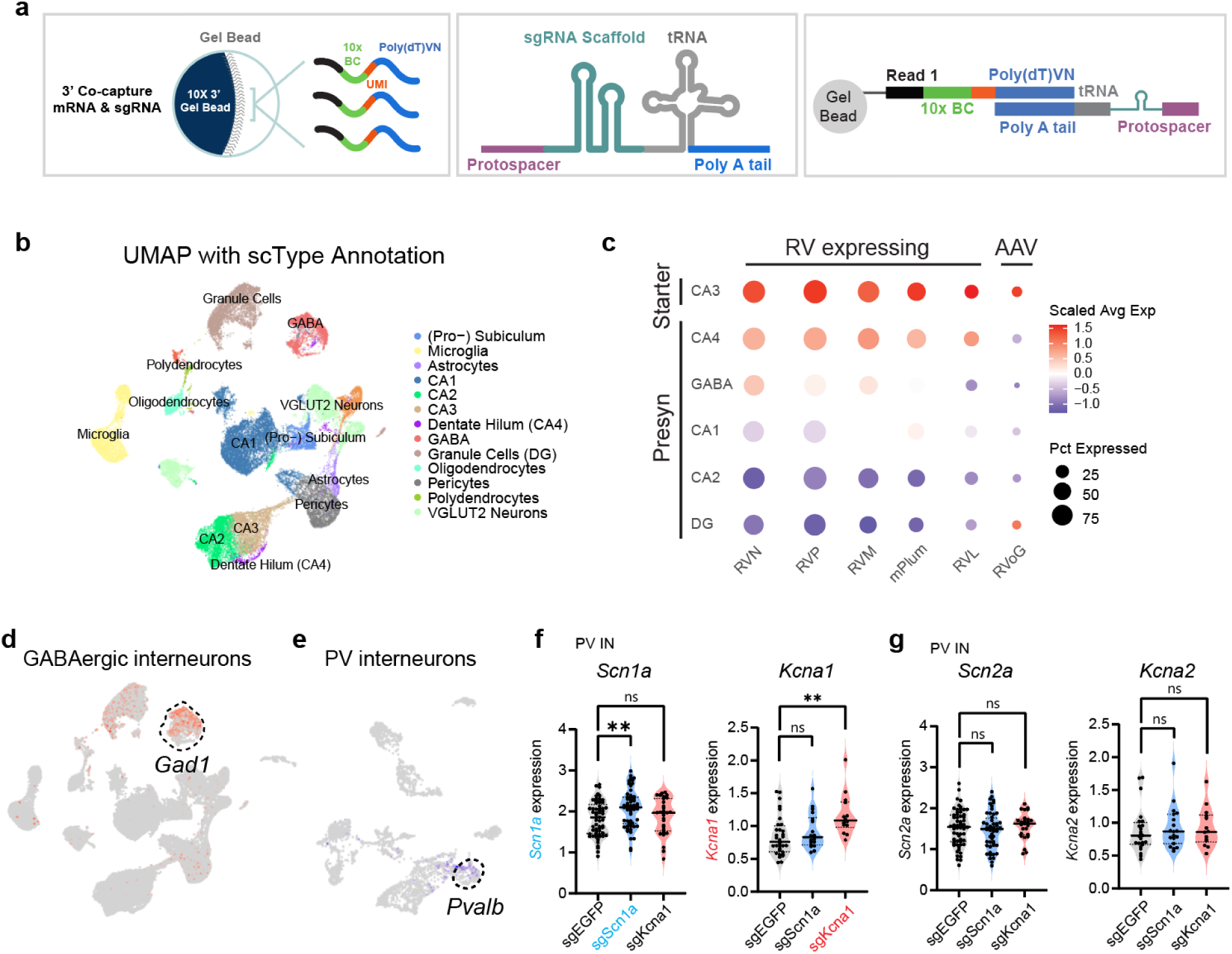
Integration of CRV-based perturbations with 3′capture single-cell sequencing for CRISPR-scRNA-seq analysis. **a)** Illustration of 3’ co-capture of mRNA and CRV-derived sgRNAs containing polyA tails. **b)** UMAP of CRV-infected nuclei identifying major hippocampal cell populations. **c)** Expression levels of rabies viral genes and AAV-mediated oG expression, in excitatory and GABAergic neuron populations. **d)** Feature plot of *Gad1***. e)** Feature plot of *Pvalb.* **f)** Violin plots showing *Scn1a* and *Kcna1* expression in PV interneurons infected with CRVs carrying different sgRNAs. **g)** Violin plots showing *Scn2a* and *Kcn1a* expression in PV interneurons infected with CRVs carrying different sgRNAs. Unpaired t-test. (**p < 0.01; ns, not significant).

## Discussion

Neural circuits represent basic functional units that underlie information processing, motor control, and cognition. Disruptions of circuit architecture are closely linked to neurological and psychiatric disorders. State-of-the-art methods, such as neuronal subtype-specific enhancers^61–63^ combined with AAV vectors and CRISPR machinery, enable gene and epigenetic editing of specific molecularly defined neuronal types. However, molecularly defined neuronal types display highly diverse projection patterns^64,65^, complicating specific neuronal components of defined circuits to produce predictable outcomes on circuit function. This problem can be thought of as adjusting an electric circuit board, which requires tuning its individual components, e.g. a specific resistor in a particular part of the circuit, rather all resistors of the same rating. Here, we introduce CRV, a novel viral CRISPR tool specifically designed for circuit- and cell-type–specific genetic manipulation. Using this approach, we achieved highly predictable changes in neuronal function based on precise gene manipulation in specific neuronal components of defined circuits, highlighting its potential for dissecting the circuit-level impact of disease-associated mutations.

Rabies virus–based tools have a long history of application in neuroscience, enabling circuit analysis not only in mammals but also in model organisms such as zebrafish. Because of its specialized life cycle, which enables trans-synaptic spread, the rabies virus serves as a powerful and reliable tool for mapping neuronal connectivity^20,23,38,66^. Recent advances have further combined rabies with high-throughput sequencing by incorporating barcode sequences into the viral genome, enabling large-scale reconstruction of neural circuits and revealing remarkable potential^34,60,67^. Nevertheless, several challenges remain. The first is toxicity: the neurons infected with the B19 vaccine strain of rabies virus can maintain electrophysiological function for only a limited period, after which they develop abnormalities and eventually degenerate^23,66,68^. The N2c strain, which is increasingly used *in vivo*, is less toxic and better tolerated but is limited by reduced transcriptional output^36,69^. In our work, we counteracted this limitation by multiplexing sgRNAs to maintain editing efficiency. Other groups have also proposed strategies to further reduce toxicity, such as deleting the L protein or fusing degradation domains^40,60,69^, thereby enabling more sustained applications. These approaches, together with the integration of long-lasting CRISPR epigenetic effectors (e.g., CRISPRoff^6^), offer promising avenues to further optimize CRV and expand its therapeutic potential. A second challenge is rabies virus production. Barcoding studies have reported that barcode sequences can mutate during viral replication^34^. Unlike AAV or lentivirus, rabies virus requires multiple rounds of self-replication during production, which risks mutation accumulation and the emergence of dominant “supercolonies”^33–35^. To ensure experimental reliability, a single-round viral recovery strategy was used, and genome sequencing–based quality control was conducted prior to use to minimize batch-to-batch variability.

An especially exciting frontier is integrating CRISPR perturbations with single-cell sequencing. This approach enables the linkage of perturbations to cell-type-specific transcriptional responses across thousands of cells in parallel. A key advantage of CRV is that its unprocessed sgRNA-tRNA transcripts retain a polyadenylated tail, making them directly compatible with standard 3’-capture platforms without requiring a switch to 5’-capture kits or specialized designs. In addition, our single-nucleus experiments revealed that rabies transcripts are highly abundant, even exceeding nuclear-localized AAV, underscoring the suitability of CRV for single-nucleus Perturb-seq. This is particularly important given that neuroscience research relies heavily on single-nucleus rather than single-cell approaches. Together, these features position CRV as a powerful and versatile platform for circuit-informed functional genomics, offering new opportunity to study how disease mutations impact defined neural circuits.

## Methods

### Plasmid construction

AAV vectors carrying sgRNAs targeting EGFP (compatible with *Staphylococcus aureus* Cas9 (SaCas9) or *Streptococcus pyogenes Cas9* (SpCas9)) were custom-designed and produced by VectorBuilder. Lentiviral plasmids expressing sgEGFP (SpCas9-compatible) were also custom-designed and produced by VectorBuilder. The pLVX-T7pol-puro construct was generated by PCR amplification of the T7 RNA polymerase (T7pol) coding sequence from pCAG-T7pol (Addgene #59926) and inserting it into pLVX-M-puro (Addgene #125839) using NEB HiFi DNA Assembly. pLenti-CMV-CAG-GFP-F2A-B19G was constructed by cloning a synthesized GFP-F2A-B19G fragment into the pLenti-CAG backbone. pSADΔG-mCherry-NLS was derived from pSADΔG-mCherry by introducing an SV40 NLS sequence via PCR and assembling with NEB HiFi DNA Assembly (NEB, E5520). All constructs were verified by whole-plasmid sequencing (Eurofins). The CRV-N2c-mCherry vector carrying a single sgRNA cassette (CRV-N2c-mCherry-1xsgEGFP) was generated by inserting a synthesized Transcription On–tRNA–sgRNA–tRNA–Off fragment (Azenta) into the RabV CVS-N2c(ΔG)-mCherry-NLS backbone (Addgene #73464). The fragment was cloned downstream of mCherry-NLS and upstream of the L gene using the NEB HiFi DNA Assembly Kit (NEB, E5520). To construct multi-copy sgRNA vectors, three overlapping PCR fragments containing homology arms were amplified from the CRV-N2c-mCherry-1xsgEGFP template and assembled using HiFi DNA Assembly to yield a 2x-sgRNA CRV vector. Additional tRNA–sgRNA cassettes were subsequently amplified from the 2x construct and randomly assembled by iterative HiFi recombination, resulting in multi-copy vectors in which sgRNAs were flanked by n+1 tRNAs. Bacterial colonies were screened using colony PCR, and correctly assembled constructs were verified by whole-plasmid sequencing (Eurofins). CRV nuclei localizing constructs were generated by inserting synthesized MCP-mCherry-NLS-2xsgEGFP-MS2, N22-mCherry-NLS-2xsgEGFP-BoxB or PCP-mCherry-NLS-2xsgEGFP-PP7 fragment into RabV CVS-N2c(ΔG)-mCherry-NLS backbone (Addgene #73464).

### Cell culture and viral transduction

N2a, P19, and 293T cell lines were obtained from the Duke Cell Culture Facility (CCF) and originally sourced from the American Type Culture Collection (ATCC). B7GG, BHK-EnvA and HEK-TVA cell lines for rabies virus production were kindly shared by Edward M. Callaway^32^.

Unless otherwise noted, cell lines were maintained at 37°C in a humidified incubator with 5% CO_2_, in Dulbecco’s Modified Eagle Medium (DMEM) supplemented with 10% fetal bovine serum (FBS) and 1% penicillin-streptomycin (P/S). P19 cells were cultured in Alpha Minimum Essential Medium (α-MEM) containing ribonucleosides and deoxyribonucleosides, supplemented with 7.5% bovine calf serum, 2.5% fetal bovine serum, and 1% P/S. All cultured cells were regularly tested for mycoplasma contamination (InvivoGen, rep-mysnc-100).

For all lentiviral transductions, cells were infected overnight in a 1:1 mixture of viral supernatant and fresh culture medium supplemented with 8 μg/mL polybrene. Transduced cells were selected with appropriate antibiotics to establish stable cell lines. Based on these, 293T cells expressing Cas9 and dEGFP and N2a cells expressing Cas9 and Tag-BFP were further transduced with lentiviruses encoding the respective effector constructs. Infected cells were isolated through FACS to enrich fluorescence positive cells. 293T7GG cells were made for rabies virus production by transducing 293T cells with lentiviruses encoding T7 polymerase and EGFP-F2A-B19G, followed by puromycin selection and FACS sorting of GFP-positive cells.

### Quantitative reverse transcription PCR (qRT-PCR)

FACS-sorted CRV-positive cells were collected, pelleted, and total RNA was extracted using the Qiagen RNeasy Mini Kit following the manufacturer’s protocol, including on-column DNase I treatment to remove genomic DNA. RNA quantity and purity were assessed by Nanodrop 2000, and 300 ng RNA was used per reaction. qRT-PCR was performed with the Takara One Step RT-qPCR kit (RR600A) on a Bio-Rad Real-Time PCR Detection System. Thermal cycling followed the kit instructions (5 min at 42 °C, initial denaturation 95 °C for 10 s, then 40 cycles of 95 °C for 5 s and 60 °C for 30 s). Reactions were run in technical triplicate. *Scn1a* and *Kcna1* expression was normalized to a *Gapdh* (See Table 1 for primer sequences). Relative expression was calculated using the 2^-ΔΔCt method, where the calibrator was RNA from mock-infected or CRV-negative sorted cells. Biological replicates are reported as mean ± s.e.m.

### Transgenic mouse breeding and maintenance

Female Rosa26-LSL-dCas9-p300 mice (dCas-p300, RRID:IMSR_JAX:033065; PMID:34341582), kindly provided by Charles Gersbach (Duke University), were crossed with male B6 Pvalb-IRES-Cre mice (PV-Cre, RRID:IMSR_JAX:017320). F1 offspring were used for stereotaxic surgery. All mice were housed in the Duke Division of Laboratory Animal Resources (DLAR) barrier facility under a 12 h light/dark cycle, with ad libitum access to food and water. Breeding pairs were maintained on LabDiet® 5058, whereas non-breeding animals were fed LabDiet® 5053.

### Virus production and purification

AAV carrying SaCas9 or SpCas9 sgRNAs were produced by VectorBuilder and purified by passing through CsCl solution. Lentivirus was produced in-house by transient transfection of 293T cells. Cells were seeded in 10 cm dishes and transfected using Lipofectamine™ 3000 (Thermo Fisher Scientific) with a three-plasmid system consisting of the expression plasmid, psPAX2 (packaging plasmid), and pMD2.G (VSV-G envelope plasmid). Culture supernatants were collected 48 and 72 hours post-transfection. Crude lentiviral supernatant was passed through a 0.45 µm PES filter, titrated, and stored directly at –80 °C. For sgRNA fidelity comparison/mutation detection, the supernatant was treated with benzonase, filtered through a 0.45 µm PES filter, and then subjected to ultracentrifugation. Filtered crude viral solution was loaded onto a 20% (w/v) sucrose cushion, SW32Ti (Beckman), 20,000 rpm, 2 h at 4 °C. Final viral preparations were resuspended in PBS, titrated, aliquoted, and stored at –80 °C until use. CRV was produced in-house by transient transfection of 293T7GG cells using the rabies genome plasmid together with helper plasmids (pcDNA-SADB19N/P/L/G for B19 strain or pCAG-N2cN/P/L/G for N2c strain). Cells were transfected with Lipofectamine 3000 and P3000 in Opti-MEM for 8 h after which the medium was replaced with complete culture medium. The following day, cells were transferred to a 35 °C incubator with 3% CO_2_. When the majority of cells exhibited red fluorescence, the culture medium was harvested, clarified by filtration, and titrated on 293T cells. Viral supernatant was aliquoted and stored at −80 °C. For sgRNA fidelity comparison and mutation detection, CRV was concentrated by ultracentrifugation, titrated, and frozen at −80 °C in parallel with lentivirus samples. For the virus used for stereotaxic surgery, crude filtered CRV supernatant was directly used to infect BHK-EnvA cells at an MOI of 0.3. After 8 h, cells were trypsinized, washed once in PBS, and replated in fresh dishes. To remove residual unpseudotyped CRV, culture dishes were washed with fresh medium before replenishing with new complete medium. EnvA-pseudotyped CRV was collected from the supernatant 3–5 days later and evaluated for titer and pseudotyping efficiency using 293TVA and 293T cells seeded in 24-well plates. Quality-checked viral supernatants were treated with benzonase, then concentrated by ultracentrifugation, followed by an additional ultracentrifugation step through a 20% sucrose cushion. Viral pellets were resuspended in PBS, titrated, and stored at −80 °C until use.

### T7E1 assay

Genomic DNA was extracted directly from cultured cells or FACS-sorted cell populations (> 100,000 cells per sample) and used as the template for PCR amplification of the target region using Q5 Hot Start High-Fidelity 2X Master Mix (NEB). Target-specific primers are listed in **Table 2**. PCR products were resolved by agarose gel electrophoresis and purified using the NucleoSpin® Gel and PCR Clean-Up Kit (Macherey-Nagel, Cat# 740609.250). DNA concentration was quantified using a NanoDrop2000 spectrophotometer. For T7E1 digestion, 200 ng of purified PCR product was subjected to a re-annealing process to allow heteroduplex formation, followed by digestion with T7 Endonuclease I (NEB, Cat# M0302) according to the manufacturer’s protocol. Digestion products were analyzed by agarose gel electrophoresis to assess the presence and efficiency of CRISPR-induced mutations.

### Western blot analysis

Cells were washed with cold PBS and counted before lysed in Cell Lysis Buffer II (with protease inhibitors) on ice for 5 minutes. Total protein concentration was measured using the Bradford assay. SDS 6×loading buffer was added to the lysates, followed by boiling at 100°C for 10 minutes. Proteins were separated by SDS–PAGE (Bio-Rad, 4561036) and transferred to nitrocellulose membranes (Bio-Rad, 1620112). Membranes were blocked with 5% BSA in TBS-T (TBS with 0.1% Tween-20) for 30 minutes at room temperature, then incubated with primary antibodies (Table 2) at 4°C overnight. After three washes with TBS-T, membranes were incubated with HRP-conjugated secondary antibodies (Table 2) at room temperature for 1 hour. Protein bands were detected using the Clarity Max ECL Substrate (Bio-Rad, 1705062S) on a Bio-Rad imaging system.

### Targeted mutation sequencing and analysis

Genomic DNA from CRV-infected cells was used as template for targeted amplification of the sgRNA protospacer region. Primers (Table 1) flanking the protospacer sequence were used for PCR with Q5 Hot Start High-Fidelity DNA Polymerase (New England Biolabs) for 10 cycles. PCR products were purified with AMPure XP beads (Beckman Coulter), and sequencing adapters were added by an additional 8–10 cycles of PCR with Q5 Hot Start to generate TruSeq Dual Index libraries. Libraries were pooled and sequenced on an Illumina NovaSeq 6000 platform with paired-end 150 bp reads (Novogene). Raw sequencing data were quality filtered using FastQC. Mutation profiles were then analyzed and visualized with CRISPResso2.

### Quantification of sgRNA mutations

Viral and plasmid indexing and NGS library preparation were performed as described previously^34^ with minor modifications. For viral samples, viral RNA/DNA was extracted from purified and concentrated CRV, AAV, or lentiviral solutions using the QIAamp Viral RNA Mini Kit (QIAGEN, 52904). 15 μL of extracted RNA/DNA was hybridized with 4 μL of 10 μM UMI-labeled oligonucleotide (e.g., sgAccuRV-UMI-F, table 1) and 4 μL of dNTPs (10 mM), then incubated at 72 °C for 4 min followed by cooling to 4 °C. For DNA plasmids, 5–25 ng of plasmid was mixed with 25 μL Q5® Hot Start High-Fidelity 2X Master Mix (NEB, M0494S), 2 μL UMI oligo, and water to 50 μL. The mixture was cycled at 98 °C for 3 min, 68 °C for 30 s, and 72 °C for 20 s. First strand synthesis was performed using Maxima H-RT (Thermo, EP0751) in a 16 μL reaction spiked into the annealed product, including Ficoll PM 400 (Sigma), RNase inhibitor, and Maxima H-RT buffer. The reaction was incubated at 42 °C for 90 min and inactivated at 85 °C for 5 min. RNA was removed by treating with RNase H (NEB, M0297S) at 37 °C for 30 min, followed by inactivation at 65 °C for 20 min. cDNA was purified using AMPure XP beads (Beckman Coulter, A63881). UMI-tagged cDNA was amplified using Q5 High-Fidelity polymerase with 10–12 cycles. Each 50 μL PCR reaction contained 25 μL purified cDNA, 0.5 μL P5 primer (table 1), 0.5 μL reverse primer targeting the sgRNA cassette (Table 1), and 25 μL Q5 Hot Start PCR mix. After bead purification, Illumina P7 index primers (Table 1) were added in a second round of amplification (10 cycles). Final PCR products were separated by agarose gel electrophoresis and gel-extracted for sequencing. Purified amplicons were sequenced on an Illumina NextSeq X platform. Mutations in sgRNA sequences were quantified by UMI collapsing and mismatch counting. Sample-specific primers used in the protocol are listed in Table 1.

### Animal preparation and stereotaxic surgery

All animal procedures complied with the NIH Guidelines for the Care and Use of Laboratory Animals and were approved by the Duke Institutional Animal Care and Use Committee (IACUC). Male and female mice (8–10 weeks old) were weighed before surgery and anesthetized with 3% isoflurane for induction. Animals were secured in a mouse adaptor on an Automated Stereotaxic Instrument (RWD, Model 71000) with a heating pad at 37 °C. Meloxicam (5 mg/kg) was administered subcutaneously for analgesia, the scalp was shaved and disinfected, and bupivacaine (0.25%, 2 mg/kg) was injected locally for additional analgesia. Throughout the procedure, anesthesia was maintained with 1.5% isoflurane delivered through a nose cone at 0.5 L/min. Injection sites were identified using coordinates from the Paxinos and Franklin Mouse Brain Atlas (2nd Edition). A burr hole (∼0.6 mm) was drilled, and viral vectors were delivered with pulled glass micropipettes using a micropump (WPI). The micropipette was left in place for 5 min after injection. The craniotomy was sealed with bone wax, and the scalp was sutured. Following recovery, mice were returned to clean cages. Post-surgical care included daily meloxicam injections on days 1 and 2. Mice exhibiting significant weight loss were euthanized according to approved protocols. For repeat surgeries, bone wax was carefully removed and the injection site cleaned before performing the procedure again as described above.

### Viral injection

Stereotaxic injections were targeted to the hippocampal CA3 at the following coordinates relative to bregma: AP −2.15 mm, ML ±2.75 mm, DV −2.5 mm. Injections were performed using pulled glass micropipettes with a total volume of 300 nL per site. For validation experiments, AAV9-Camk2a-TVAoG was injected at 300 nL (1×10¹³ GC/mL). For experiments involving sEPSC/sIPSC recordings, immunofluorescence, and single-nucleus RNA sequencing, mice received a first injection of AAV9-Camk2a-TVAoG (300 nL at 2×10¹² GC/mL), followed by a second injection of CRV (300 nL at 5×10⁸ TU/mL). To identify CRV-infected PV neurons, mice were first injected with AAV9-Camk2a-TVAoG (300 nL at 2×10¹² GC/mL), followed by a second injection of CRV (300 nL at 5×10⁸ TU/mL) mixed with AAV PHP.Eb-FLEX-tdTomato (2×10¹² GC/mL).

### Histological analysis

Mice were anesthetized with isoflurane and perfused transcardially with cold PBS followed by 4% paraformaldehyde (PFA). Brains were dissected, fixed in 4% PFA overnight at 4 °C, and transferred to 30% sucrose in PBS 1-2 days for cryoprotection. Tissues were embedded in OCT & Sucrose (30%) compound (3:2 v/v) and sectioned on a cryostat. For free-floating immunostaining, sections were permeabilized with 0.4% Triton X-100 in PBS for 5 min and blocked with 3% BSA in PBS for 1 h at room temperature. Sections were incubated with primary antibodies (Table 2) at 4 °C overnight, washed three times with PBST (PBS + 0.1% Tween-20), and then incubated with secondary antibodies for 2 h at room temperature. After washing, sections were counterstained with DAPI and mounted. Images were acquired using a Zeiss LSM 880 confocal microscope.

### RNAscope *in situ* hybridization (ISH)

To assess whether CRV infection increases nuclear rabies RNA, cells infected with different CRV constructs were first FACS-sorted and then processed for nuclei isolation. Sorted cells were homogenized on ice in nuclei isolation buffer (0.32 M sucrose, 5 mM CaCl_2_, 3 mM MgAc_2_, 0.1 mM EDTA, 10 mM Tris-HCl pH 8.0, 1 mM DTT, 0.10% NP-40, and 1× protease inhibitors) using a Dounce homogenizer. The homogenate was layered carefully along the wall of ultracentrifuge tubes (Beckman, 355631) containing a pre-chilled sucrose cushion (1.8 M sucrose, 3 mM MgAc_2_, 10 mM Tris-HCl pH 8.0, 1 mM DTT), and centrifuged at 20,000 rpm for 1 h at 4 °C in a SW 32 Ti rotor (Beckman). The nuclear pellet was collected, washed, and resuspended in nuclei isolation wash buffer (3 mM MgAc_2_, 10 mM Tris-HCl pH 7.4, 10 mM NaCl, 1% BSA, 0.10% Tween-20), followed by incubation on ice for 20 min. Nuclei were gently resuspended, mounted onto poly-L-lysine–coated glass slides, and air-dried. RNAscope ISH was carried out with probes targeting rabies viral genes (RVG and RVM) using the manufacturer’s protocol (ACD Bio). After hybridization and signal amplification, nuclei were coverslipped and imaged with a Zeiss LSM 880 confocal microscope. The number of rabies RNA foci per nucleus was quantified from confocal images.

### Electrophysiological recordings

After urethane anesthesia (1.5–2.0 g/kg, i.p.), mice were transcardially perfused with 20 ml ice-cold (1–4 °C) dissection solution (240 mM sucrose, 25 mM NaHCO₃, 2.5 mM KCl, 1.25 mM NaH_2_PO₄, 0.5 mM CaCl_2_, and 3.5 mM MgCl_2_). Brains were rapidly removed and immersed in oxygenated (95% O_2_ and 5% CO_2_) ice-cold dissection solution. Transverse hippocampal slices (350 μm) were prepared using a vibrating microtome (VT1200S; Leica). Slices were incubated at 32 °C for 30 min in ACSF (NaCl 117 mM, KCl 3.6 mM, NaH_2_PO₄·2H_2_O 1.2 mM, CaCl_2_·2H_2_O 2.5 mM, MgCl_2_·6H_2_O 1.2 mM, NaHCO₃ 25 mM, D-glucose 11 mM, and sodium pyruvate 2 mM, pH 7.4) equilibrated with 95% O_2_ and 5% CO_2_.

mPlum (red channel) and EGFP (green channel) double-positive neurons in the hippocampus CA3 were identified under epifluorescence illumination (BX51WIF; Olympus; Lumen 200 Fluorescence Illumination System, PRIOR). Whole-cell voltage-clamp recordings were performed using an Axopatch 200B amplifier (Molecular Devices, San Jose, CA), filtered at 2 kHz and digitized at 5 kHz. Patch pipettes had a resistance of 6–10 MΩ. For sEPSC recordings, cells were held at –70 mV and pipettes were filled with internal solution containing 135 mM K-gluconate, 0.5 mM CaCl_2_, 2 mM MgCl_2_, 2.5 mM EGTA, 5 mM HEPES, and 5 mM Mg-ATP (pH 7.3). For sIPSC recordings, cells were held at 0 mV and pipettes contained 110 mM CsSO₄, 0.5 mM CaCl_2_, 2 mM MgCl_2_, 5 mM TEA, 5 mM EGTA, 5 mM HEPES, and 5 mM Mg-ATP (pH 7.3). Data were acquired using pClamp 10.7 software (Molecular Devices) and analyzed with MiniAnalysis (ver. 6.0.7; Synaptosoft, Decatur, GA). Thresholds for sEPSC and sIPSC detection were set at 7 pA and 9 pA, respectively and events were further verified by manual inspection of waveform morphology. Excitatory currents were pharmacologically isolated by adding 10 µM picrotoxin to the bath solution. During recordings, slices were perfused with ACSF (2–4 ml/min) saturated with 95% O_2_ and 5% CO_2_ at room temperature.

### Flow Cytometry and Fluorescence-Activated Cell Sorting (FACS)

For flow cytometric analysis, cells were dissociated using trypsin, washed with PBS, and passed through a 40 µm cell strainer. Flow cytometry was performed using a BD LSRFortessa™ Cell Analyzer to assess viral infection efficiency (mCherry expression) and editing efficiency (loss of EGFP fluorescence). To generate stable cell lines expressing fluorescent proteins, cells were transduced with lentiviral vectors and subsequently sorted using a Sony MA900 cell sorter equipped with a 100 µm microfluidic chip. Single-cell populations were gated based on forward and side scatter profiles, and sorted into fluorescence-positive populations. Sorted cells were expanded in culture and re-analyzed by fluorescence microscopy to confirm stable expression. Nuclei sorting procedures are described in the Nuclei Isolation and Single-Cell RNA Sequencing section.

### Nuclei isolation and single-cell RNA sequencing

Hippocampal tissue was rapidly frozen in 2-methylbutane pre-cooled on dry ice and stored at – 80 °C until processing. For nuclear isolation, tissue was incubated with freshly prepared, ice-cold nuclei extraction buffer^70^, followed by mechanical dissociation using a Dounce homogenizer on ice until no visible tissue pieces, as described in the RNAscope ISH section. After isolation, nuclei were stained with DAPI (final concentration: 1 µg/mL) on ice for 10 min, and DAPI⁺/H2B-mPlum⁺ nuclei were sorted using a Sony MA900 cell sorter equipped with a 100 µm microfluidic chip. Sorted nuclei were before loading. Single-nucleus RNA-seq libraries were prepared using the 10x Genomics Chromium Single Cell 3’ v4 kit. Approximately 20,000 nuclei per well were loaded with one well containing nuclei from the sgKcna1 group, and another containing a mixture of nuclei from the sgEGFP and sgScn1a groups. cDNA generated from each well was divided, with one portion used to construct the standard gene expression library and the other portion subjected to targeted enrichment of sgRNA transcripts to identify sgRNA-expressing nuclei. Libraries were sequenced on an Illumina NovaSeq X platform with paired-end 150 bp reads.

## Data and materials availability

All data supporting the findings of this study are available within the paper and its supplementary materials. Key CRV plasmids will be deposited at Addgene.

## Acknowledgments

We thank all members of the Dmitry Velmeshev lab for their helpful advice and discussions. We thank the Duke Core Facilities for assistance with animal work, flow cytometry, and imaging. We are grateful to Dr. Z. Josh Huang and Baoxia Han for their guidance on transgenic mouse genotyping and breeding. This study is supported by the NIH R00 grant R00MH121534 and DP1 grant DP1DA063507, Duke University’s Whitehead Scholarship, and a generous donation by David Dolby to D.V. D.S. is supported by the NIH 1DP2MH140149, the Klingenstein-Simons Fellowship Award in Neuroscience, the Whitehall Foundation, and the Ruth K. Broad Foundation for Biomedical Research. Z.Z. is supported by a postdoctoral fellowship from the Ruth K. Broad Biomedical Research Foundation.

## Author contributions

Conceptualization: ZZ, DV

Methodology: ZZ, JX, EM, JM, DS, CG, RJ, DV

Investigation: ZZ, JX, EM, SD, LH, DA, EP, AD, KSK

Visualization: ZZ, JX, DV

Funding acquisition: DV

Project administration: DV

Supervision: DV

Writing – original draft: ZZ, DV

Writing – review & editing: ZZ, JX, EM, SD, LH, DA, EP, AD, JM, DS, CG, RJ, DV

## Supplementary Materials

### Supplementary figure legends

**Figure s1.**
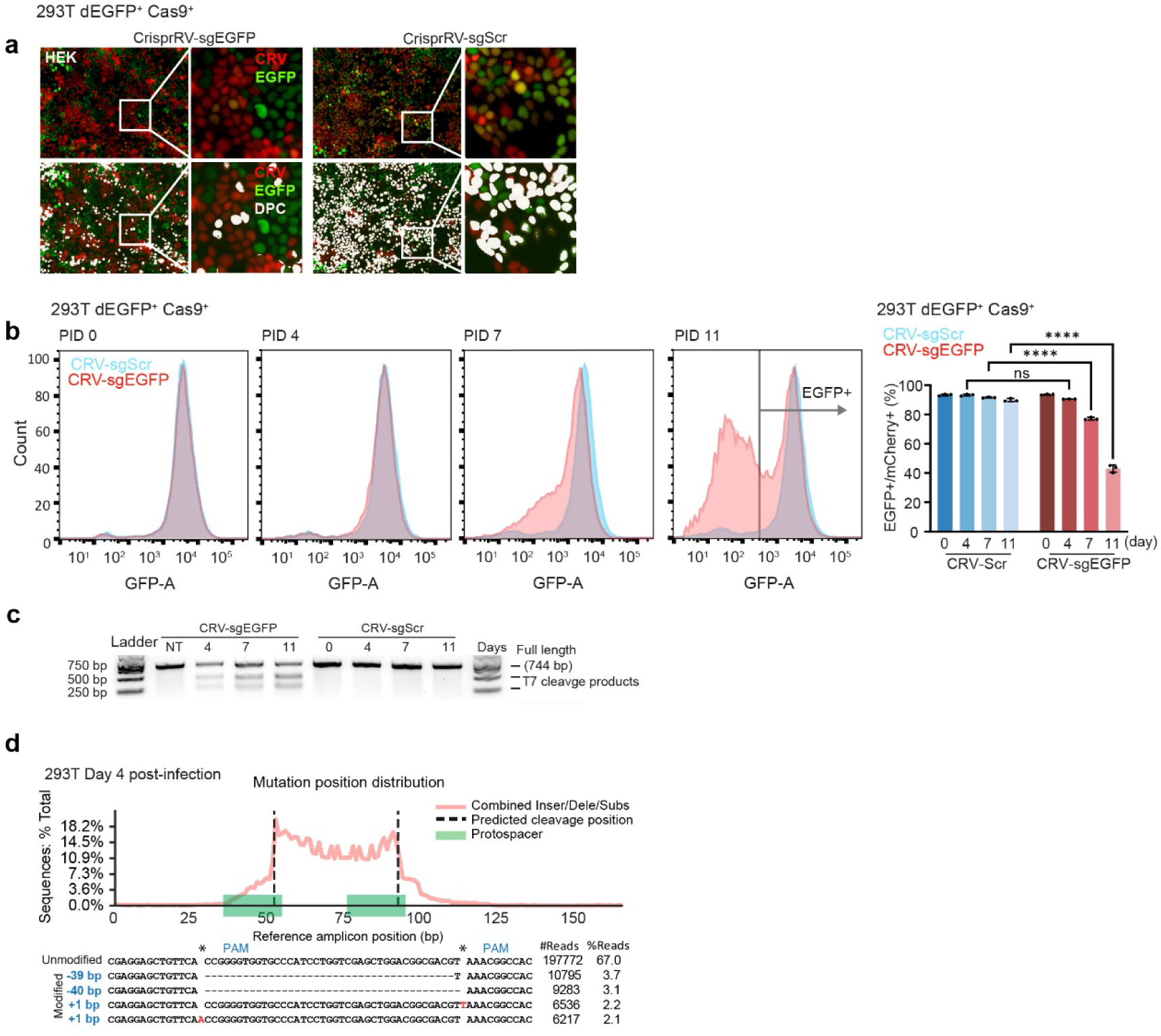
Validation of CRV. **a)** 293T-dEGFP-Cas9 cells infected with CRV expressing nuclear-localized mCherry and carrying either an *EGFP*-targeting sgRNA or a scrambled control. Double-positive cells (DPCs) are highlighted in white. **b)** Flow cytometry analysis of *EGFP* expression in infected cells was performed from day 0 to day 11. Data are presented as mean ± s.d. Statistical significance was assessed by two-way ANOVA with multiple comparisons (****p < 0.0001; ns, not significant). **c)** T7E1 assay on the EGFP locus amplified 0-11 days after infection at MOI = 1. **d)** CRISPResso2 analysis of mutation position distribution at the EGFP target locus in 293T cells from c) infected with CRV for 4 days.

**Figure s2.**
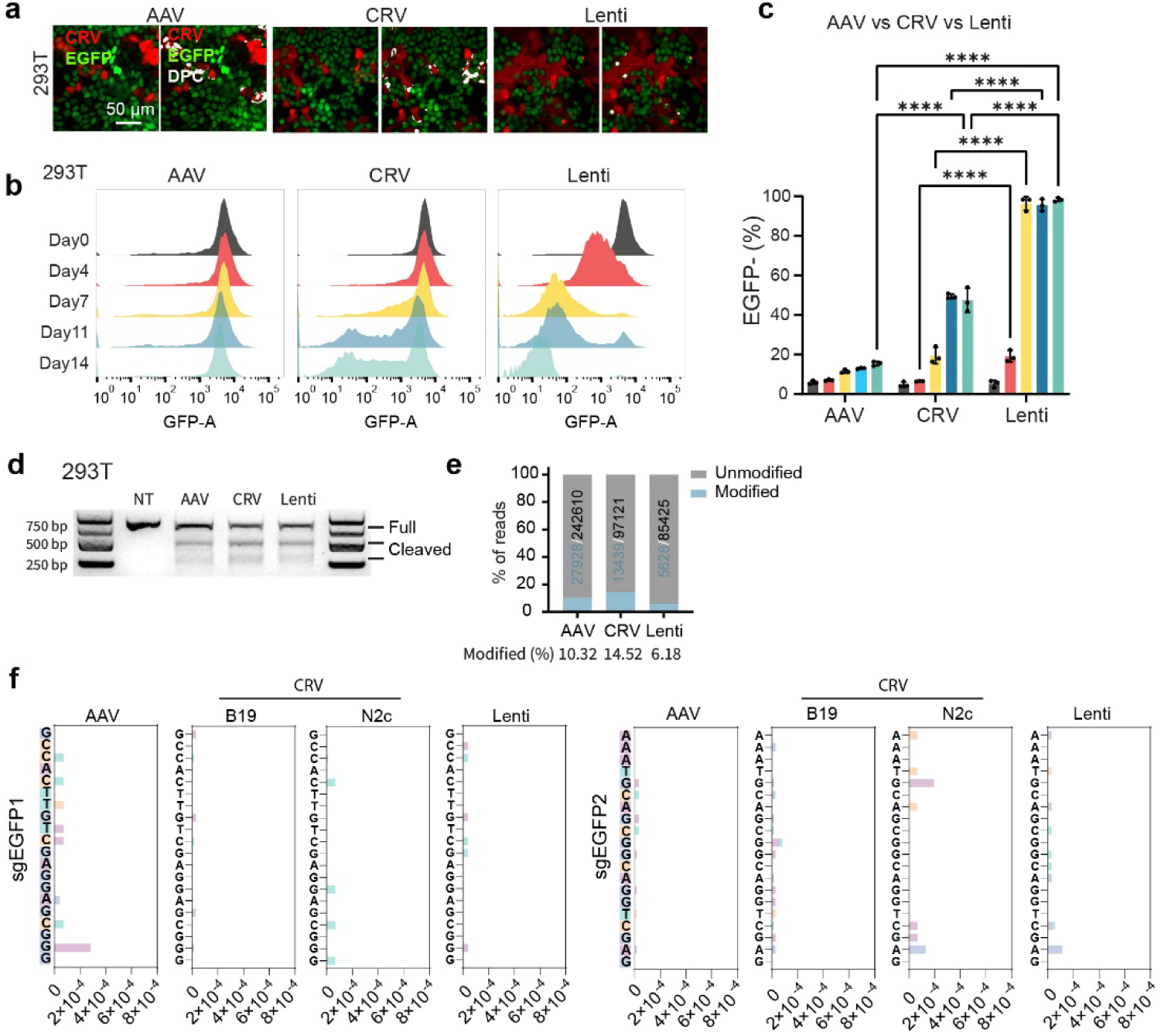
Comparison of efficiency and reliability between CRV, AAV, and lentiviral sgRNA delivery systems. **a)** AAV, CRV, and lentivirus expressing mCherry together with 2 identical sgRNAs targeting EGFP. At day 11 post-infection, red (mCherry⁺, infected) 293T-dEGFP-Cas9 cells losing green fluorescence indicate successful targeting. DPCs are highlighted in white. **b-c)** Flow cytometry analysis of EGFP expression from day 0-14 post AAV, CRV and lentivirus infected red cells (mCherry⁺). Data are presented as mean ± s.d. Statistical significance was assessed by two-way ANOVA with multiple comparisons (**p < 0.0001; ns, not significant). **d)** T7E1 assay on the EGFP locus amplified from genomic DNA of mCherry⁺ cells, FACS-sorted 4 days after infection at MOI = 0.3 (CRV, Lenti) or ∼100,000 (AAV). **e)** Bar plot showing mutation frequency by NGS of adapter-ligated PCR products from d). **f)** Mutation frequency and types for sgEGFP1 and sgEGFP2 in plasmids used for viral production.

**Figure s3.**
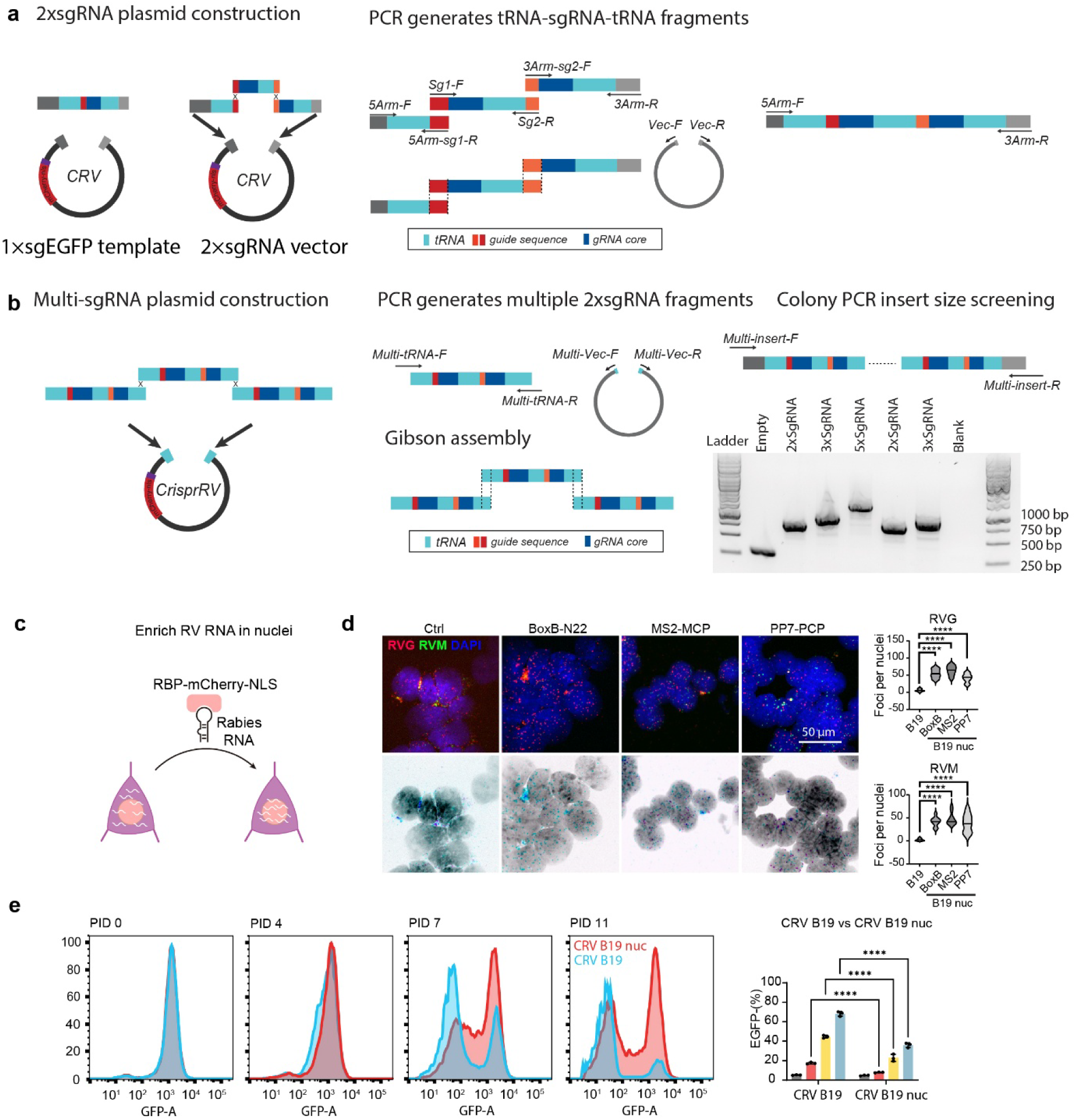
Construction and validation of multiplexed tRNA–sgRNA and nuclei-localized CRV vectors. **a)** Schematic illustration of CRV constructs carrying 2× sgRNAs. **b)** Schematic illustration of CRV constructs carrying multiple sgRNAs and colony PCR screening of positive bacterial clones. **c)** Schematic illustration of CRV-B19 nuc infected cell, in which fusion of RNA binding proteins (RBP) with nuclear-localized fluorescent proteins enables capture of RNAs containing BoxB, MS2, or PP7 structures. **d)** RNAscope detection of rabies viral RNA distribution, with violin plots showing the number of nuclear rabies foci per nuclei. **e)** CRV B19 (MS2–MCP) infection of 293T-dEGFP-Cas9 cells carrying sgRNAs targeting EGFP. Flow cytometry analysis of EGFP expression from day 0 to day 11 & quantification of EGFP negative cells.

**Figure s4.**
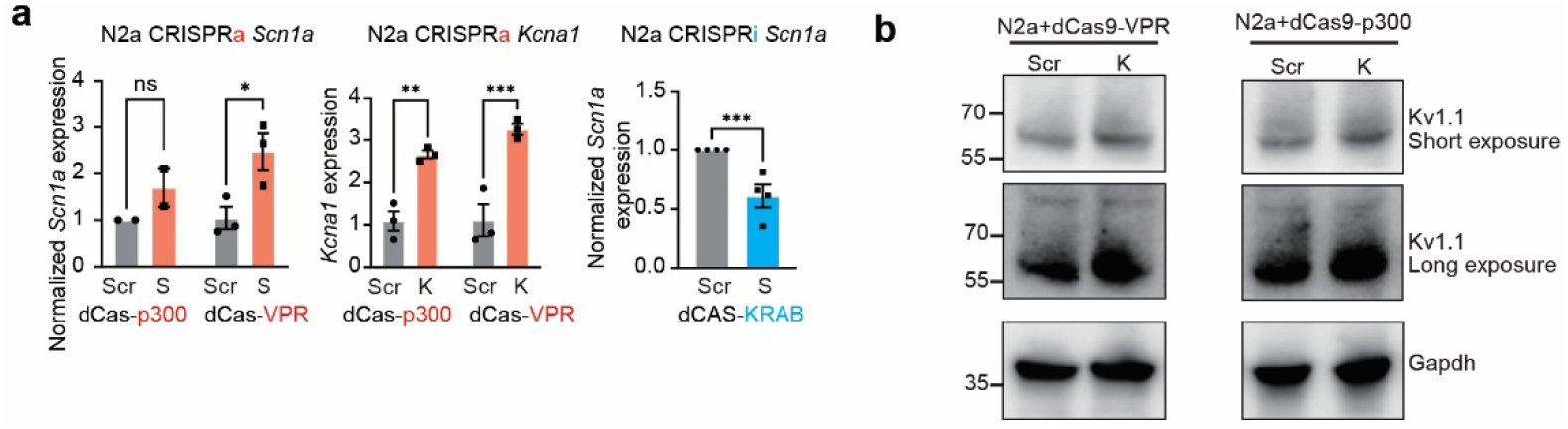
CRV targets endogenous genes. **a)** Relative mRNA expression of Scn1a and Kcna1, determined by qRT-PCR, in cells expressing dCas9 and infected with CRVs carrying the indicated sgRNAs. **b)** Protein expression levels of Kv1.1 determined by immunoblotting, in cells expressing dCas9 and infected with CRV carrying the indicated sgRNAs. (Scr, scrambled control; K, sgKcna1; S, sgScn1a). Unpaired t-test. Data are presented as mean ± s.e.m.

**Figure s5.**
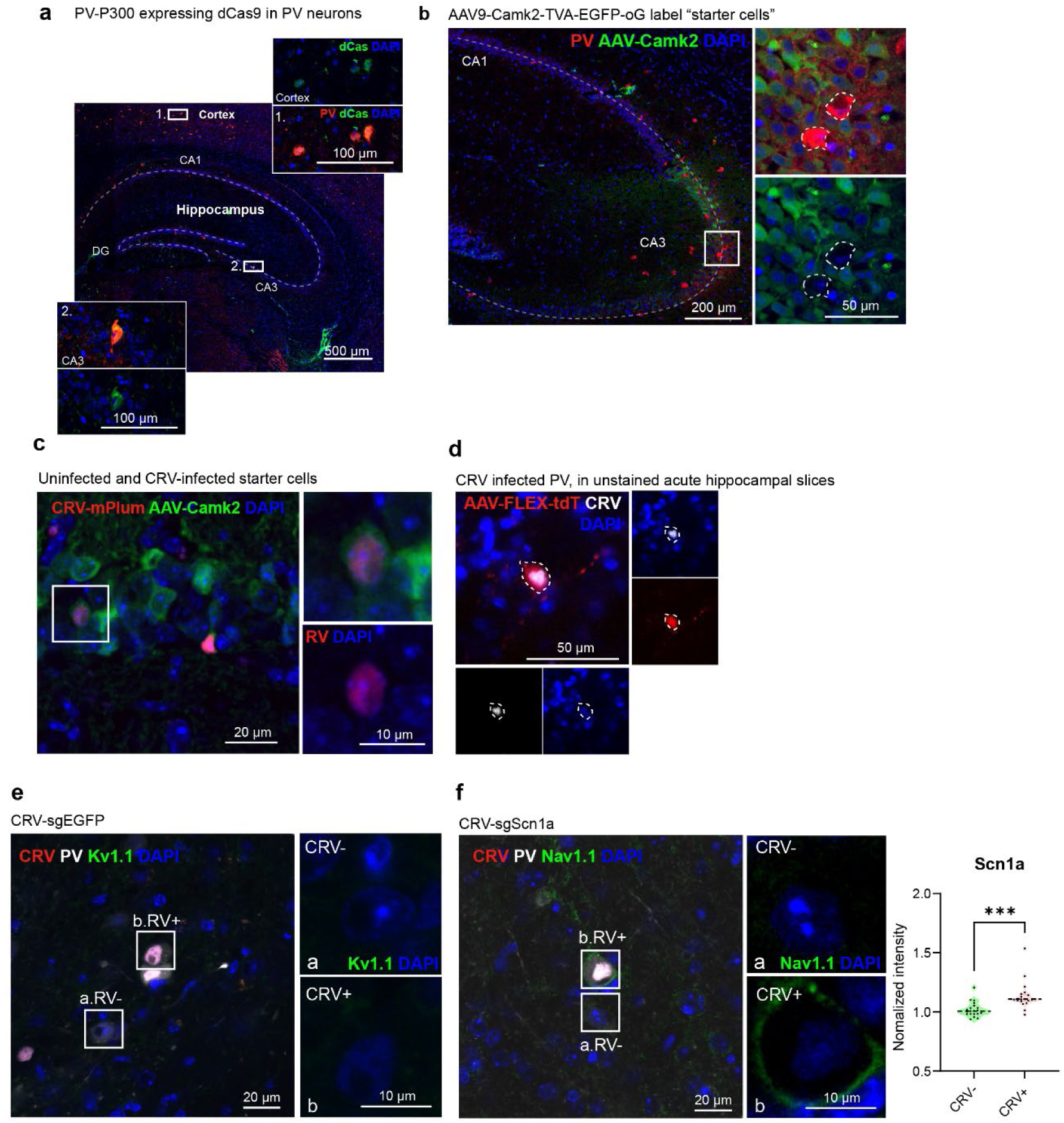
CRV-mediated circuit specific targeting of PV neuron. **a)** Immunofluorescence (IF) detection of dCas9 expression in PV–p300 hippocampal slices. Scale bar, 100 µm. **b)** IF detection of AAV-expressed EGFP and PV neuron marker Parvalbumin. **c)** IF detection of AAV-expressed EGFP and CRV-expressed mPlum, showing starter cells and CRV-infected starter cells. **d)** Unstained acute hippocampal slices showing AAV-FLEX-tdTomato labeled PV neurons and CRV-expressed H2B-mPlum, illustrating CRV infection of PV neurons. **e)** Immunofluorescence confocal imaging of Kv1.1 expression in PV neurons shows increased Kv1.1 signal in the CRV-sgEGFP–infected neuron compared with an adjacent uninfected PV neuron. **f)** Immunofluorescence analysis of Nav1.1 expression in PV neurons infected with CRV-sgScn1a and an adjacent uninfected PV neuron. Unpaired t-test (*p < 0.001).

**Figure s6.**
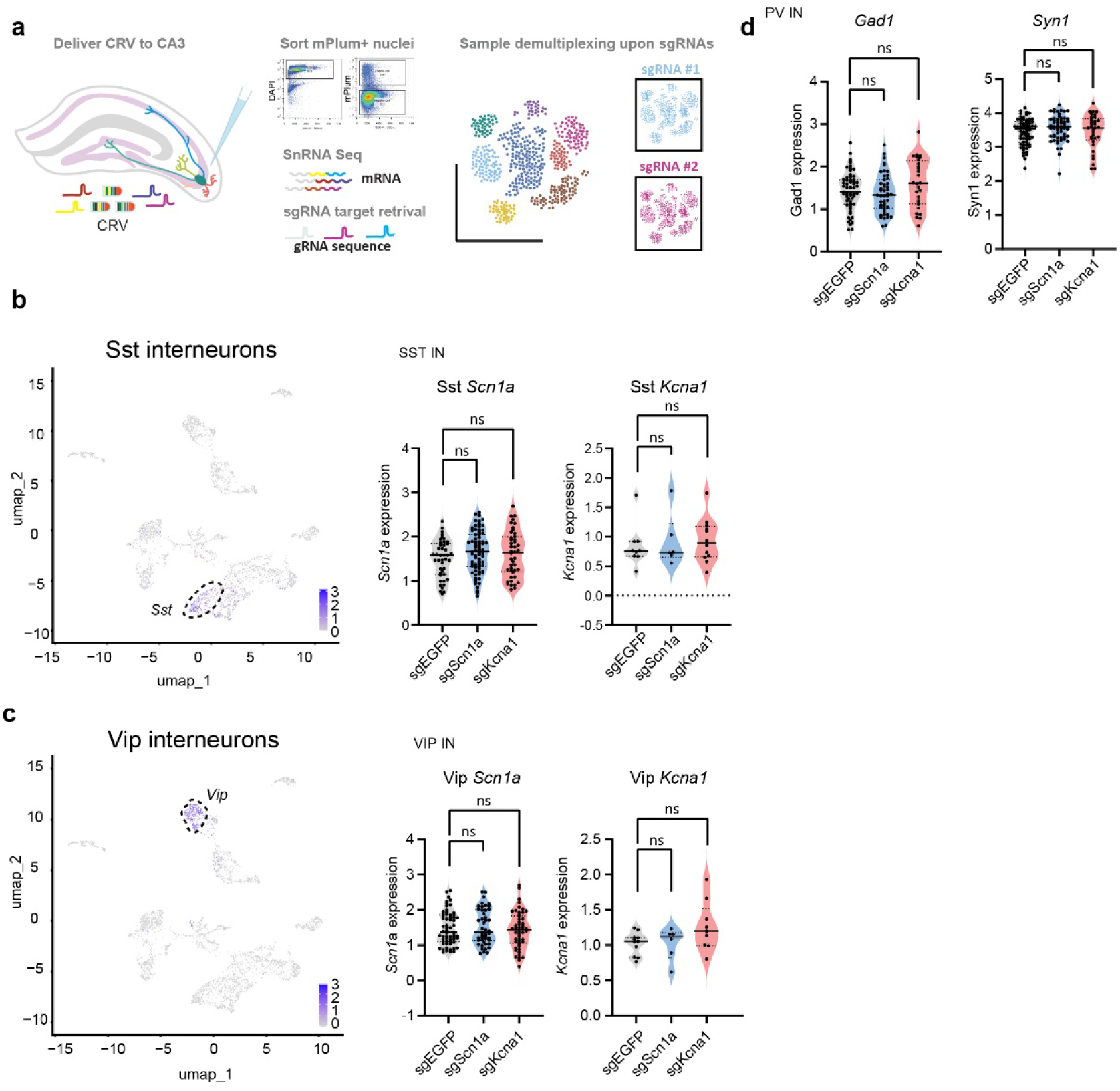
Integration of CRV-based perturbations with 3′capture single-cell sequencing for CRISPR-scRNA-seq analysis. **a)** Illustration of CRV infection, nuclei isolation, and protospacer-based sample demultiplexing post sequencing. Sample demultiplexing was achieved using protospacer sequence information embedded in the sgRNA design, allowing accurate assignment of nuclei to their respective perturbations. **b)** Violin plots showing expression levels of housekeeping genes in PV neurons infected with CRVs carrying different sgRNAs (ns, not significant). **c)** Feature plot of Sst and violin plots showing *Scn1a* and *Kcna1* expression in Sst interneurons infected with CRVs carrying different sgRNAs. **d)** Feature plot of Vip and violin plots showing *Scn1a* and *Kcna1* expression in Vip interneurons infected with CRVs carrying different sgRNAs. Unpaired t-test. (**p < 0.01; ns, not significant).

